# Using Functional Genomic Data in Monocytes/Macrophages and Genotyping to Nominate Disease-Driving Single Nucleotide Polymorphisms and Target Genes in Juvenile Idiopathic Arthritis

**DOI:** 10.1101/2024.08.19.608312

**Authors:** Emma K. Haley, Gilad Barshad, Adam He, Edward Rice, Marc Sudman, Susan D. Thompson, Elizabeth A. Crinzi, Kaiyu Jiang, Charles G. Danko, James N. Jarvis

## Abstract

**Introduction:** GWAS have identified multiple regions that confer risk for juvenile idiopathic arthritis (JIA). However, identifying the single nucleotide polymorphisms (SNPs) that drive disease risk is impeded by the SNPs’ that identify risk loci being in linkage disequilibrium (LD) with hundreds of other SNPs. Since the causal SNPs remain unknown, it is difficult to identify target genes and use genetic information to inform patient care. We used genotyping and functional data in primary human monocytes/macrophages to nominate disease-driving SNPs on JIA risk haplotypes and identify their likely target genes.

**Methods:** We identified JIA risk haplotypes using Immunochip data from Hinks et al (Nature Gen 2013) and the meta-analysis from McIntosh et al (Arthritis Rheum 2017). We used genotyping data from 3,939 children with JIA and 14,412 healthy controls to identify SNPs that: (1) were situated within open chromatin in multiple immune cell types and (2) were more common in children with JIA than the controls (p< 0.05). We intersected the chosen SNPs (n=846) with regions of bi-directional transcription initiation characteristic of non-coding regulatory regions detected using dREG to analyze GRO-seq data. Finally, we used MicroC data to identify gene promoters interacting with the regulatory regions harboring the candidate causal SNPs.

**Results:** We identified 190 SNPs that overlap with dREG peaks in monocytes and126 SNPs that overlap with dREG peaks in macrophages. Of these SNPs, 101 were situated within dREG peaks in both monocytes and macrophages, suggesting that these SNPs exert their effects independent of the cellular activation state. MicroC data in monocytes identified 20 genes/transcripts whose promoters interact with the enhancers harboring the SNPs of interest.

**Conclusion:** SNPs in JIA risk regions that are candidate causal variants can be further screened using functional data such as GRO-seq. This process identifies a finite number of candidate causal SNPs, the majority of which are likely to exert their biological effects independent of cellular activation state in monocytes. Three-dimensional chromatin data generated with MicroC identifies genes likely to be influenced by these SNPs. These studies demonstrate the importance of investigations into the role of innate immunity in JIA.

## Introduction

Juvenile idiopathic arthritis (JIA) is one of the most common chronic disease conditions affecting children in North America(1, 2). While the pathogenesis of JIA remains unknown, genetic factors are known to contribute to disease risk. For example, in a study using the Utah Population Registry, Prahalad et al demonstrated that the relative risk for developing JIA in siblings of affected children was nearly 12 times that of the general population (OR =11.6; confidence intervals 4.9–27.5; p<3×10^−8^), and that for first cousins was nearly 6 times (OR =5.8; confidence intervals 2.5–13.8; p<6×10^−5^)(3). More recently, genome-wide association studies (GWAS) and genetic fine mapping studies(4, 5) have identified 23 regions outside of the major histocompatibility complex (MHC) that confer disease risk. It is likely that additional risk regions will be identified as larger patient cohorts are assembled.

Despite their utility, GWAS cannot and were not designed to answer key questions about genetic risk. Because of the phenomenon of linkage disequilibrium (LD), GWAS are unable to identify the actual variants that exert the biological effects that drive disease risk. If one considers only the non-MHC regions identified in the Hinks Immunochip and McIntosh meta-analysis papers for example, there are >13,000 SNP in LD with those that tag the genetic risk loci. Furthermore, GWAS are unable to identify the genes whose functions/expression levels are influenced by the disease-driving variants. This problem is made particularly complex because: (1) genetic risk is likely to be exerted mainly through non-coding, regulatory elements(6-8), a phenomenon common across autoimmune diseases(9); (2) regulatory elements may not influence the most proximal gene (in terms of linear genomic distance) to either the tag or causal SNP, and, thus, the 3D structure of the genome must be considered(10). Finally, GWAS do not identify the cells or tissues affected by the disease-driving variants, a crucial question that must be answered to deliver the precision medicine approaches that the field has long sought. We have published data showing that disease-driving variants in JIA are likely to influence multiple immune cell type(5, 8, 10), findings that are corroborated by the Genotype-Tissue Expression (GTEx) project(11), where single variants can be shown to influence gene expression in a range of functionally different tissues.

In this paper, we demonstrate that using existing genotyping data and identifying important functional features in individual cell types, we can considerably reduce the list of potential disease-driving variants on JIA risk haplotypes. Furthermore, we show that using 3D chromatin data can assist in identifying the likely target genes of the disease-driving variants, and, thus, the immunologic processes impacted. We chose monocytes/macrophages for these analyses because, although this cell type is known to contribute to JIA pathogenesis(12, 13), it is still under-studied, leaving an important gap in our understanding of the disease pathogenesis.

## Methods

### Genotyping data sets

We queried raw data from genome-wide association studies and genetic fine mapping studies as previously reported in(4, 5). From McIntosh et al. three cohorts comprising 2,751 patients with oligoarticular or RF-negative polyarticular JIA were genotyped using the Affymetrix Genome-Wide SNP Array 6.0 or the Illumina HumanCoreExome-12+ Array. Overall, 15,886 local and out-of-study controls, typed on these platforms or the Illumina HumanOmni2.5, were used for association analyses.

From Hinks et al. Immunochip array was used to analyze 2,816 individuals with juvenile idiopathic arthritis (JIA), comprising the most common subtypes (oligoarticular and rheumatoid factor-negative polyarticular JIA), and 13,056 controls. Combined, these studies include genotyping data from 3,939 children with oligoarticular/polyarticular (RF-negative) JIA and 14,412 healthy controls. Using r^2^ >0.50, we identified all SNPs on those haplotypes listed in **Table 1** that were in linkage disequilibrium (LD) with the tag SNPs. We further filtered these SNPs using the QC analysis methods described in(4, 5) (e.g., presence of Hardy Weinberg equilibrium, minor allele frequency, missingness), and identified those that were significantly more common in patients than in controls (p <0.05). Finally, from this list, we identified SNPs that were situated within regions of open chromatin and within H3K4me1/H3K27ac peaks in human neutrophils and CD4+ T cells(7). These filtering processes provided us with a list of 846 candidate SNPs in **Supplemental Table 1**.

**Table 1:** Positional information of the 23 JIA-risk single nucleotide polymorphisms and the associated haplotypes.

| <u>Locus Name</u> | <u>SNP</u> | <u>Hg38 Haplotype</u> |
| --- | --- | --- |
| PTPN22 | rs6679677 | chr1:113,761,186-113,834,946 |
| STAT4 | rs10174238 | chr2:191,079,016-191,108,308 |
| PTPN2 | rs2847293 | chr18:12,774,327-12,809,341 |
| ANKRD55 | rs71624119 | chr5:56,141,024-56,146,422 |
| IL2-IL21 | rs1479924 | chr4:122,151,854-122,619,603 |
| TYK2 | rs34536443 | chr19:10,317,045-10,381,598 |
| IL2RA | rs7909519 | chr10:6,028,313-6,055,320 |
| SH2B3-ATXN2 | rs3184504 | chr12:111,395,984-111,645,358 |
| ERAP2-LNPEP | rs27290 | chr5:96,884,383-97,038,046 |
| UBE2L3 | rs2266959 | chr22:21,556,931-21,628,971 |
| C5orf56/IRF1 | rs4705862 | chr5:132,477,527-132,496,822 |
| RUNX1 | rs9979383 | chr21:35,323,611-35,365,944 |
| IL2RB | rs2284033 | chr22:37,135,077-37,141,474 |
| ATP8B2/IL6R | rs11265608 | chr1:154,319,242-154,406,893 |
| ZFP36L1 | rs12434551 | chr14:68,784,174-68-794,755 |
| JAK1 | rs10889504 | chr1:64,924,820-64,975,081 |
| PRR9/LOR | rs873234 | chr1:153,249,128-153,270,022 |
| PTH1R | rs1138518 | chr3:46,889,988-46,932,682 |
| ILDR1/CD86 | rs111700762 | chr3:122,023,892-122,102,059 |
| AHI1/LINC00271 | rs9321502 | chr6:135,303,673-35,371,709 |
| HBP1 | rs111865019 | chr7:107,156,092-107,390,877 |
| WDFY4 | rs1904603 | chr10:48,776,493-48,805,795 |
| RNF215 | rs57533109 | chr22:30,289,571-30,403,731 |

chr chromosome, rs reference SNP, SNP single nucleotide polymorphism

### Assessing functional regions in human monocyte-macrophages

We focused on the JIA risk regions as identified in the Hinks genetic fine mapping study(5) and McIntosh meta-analysis(4). To identify linkage disequilibrium blocks associated with the risk SNPs, we used the Single Nucleotide Polymorphism Annotator (SNiPA) (21), available publicly at https://snipa.helmholtz-muenchen.de/snipa3/. We queried European populations, i.e., the populations represented in the referenced genetic studies, using GRCh37/Ensembl 87. Although our initial set of n=846 SNPs was generated using r^2^ = 0.5, we focused this study on the regions identified using r^2^ = 0.80. We subsequently converted GRCh37/hg19 coordinates to GRCh38/hg38 using the liftover tool to map relevant chromatin data. These regions are shown in **Table 1**.

### Assessing functional regions in monocyte-macrophages

We queried global run-on sequencing (GRO-seq) data using the dREG analysis pipeline(13) to identify non-coding regulatory elements in publicly available data from resting CD14+ monocytes and macrophages activated with Kdo2-Lipid-A (KLA), a lipopolysaccharide derived from E. coli ([GSM1183908 and GSM1183912] (14). As described in our previous paper we used the intersect command in BedTools to identify GRO-seq peaks within the JIA LD blocks and dREG analysis to identify functional regions(8).

### Micro-C to Identify Interacting Regions in Human Monocytes

#### Monocyte Isolation

The isolation of CD14+ monocytes human blood was done in compliance with Cornell University IRB guidelines. We received peripheral blood samples (60–100 mL) from healthy adult humans. Subjects (two male and three female subjects) gave informed consent for inclusion in this study. Blood was collected in purple top EDTA tubes and maintained at 4 °C overnight before processing. Blood was diluted 50:50 with PBS and peripheral blood mononuclear cells (PBMCs) were separated by applying 20 mL of blood:PBS mixture over 20 mL Ficoll-Pacque and centrifuging (750×g) for 30 min at 20 °C to create a gradient. Cells from the PBMC layer of the Ficoll gradient were collected and washed three times in ice-cold PBS supplemented with 0.5% BSA. If red blood cell contamination occurred, the cell pellet was treated with ACK red blood cell lysis buffer. CD14 + Monocytes were isolated using CD14 microbeads (Miltenyi Biotech, 130-050-201) according to manufacturer’s protocol: A total of 10^8^ cells were resuspended in MACS binding buffer and bound to microbeads (20 µL of microbeads/10^7^ cells) for 15 min at 4 °C. Cells were washed with 1-2 mL of PBS/BSA solution, resuspended in 500 µL of binding buffer, and passed over a MACS LS column (Miltenyi Biotech, 130-042-401) on a magnet at 4 °C. The MACS LS column was washed three times with 2 mL PBS/BSA solution, before cells were eluted off the magnet. Cells were counted using a hemocytometer, resuspended in RPMI-1600, and incubated at 37 °C for one hour before further processing.

#### Micro-C

Cells were crosslinked with 1 ml per million cells of 1% formaldehyde for 10 minutes at room temperature and quenched by 0.25 M Glycine for 5 min. After spin-down for 5 minutes at 300Xg at 4 °C, cells were washed at a density of 1 ml per million cells in ice cold PBS. Cells were crosslinked a second time, with 1 ml per 4 million cells of 3 mM disuccinimidyl glutarate (DSG) (ThermoFisher Scientific, 20593) for 40 min at room temperature and quenched by 0.4 M Glycine for 5 min. Following two washes with ice cold PBS, cells were flash-frozen and kept at -80°C until further use.

For MNase digestion, cells were thawed on ice for 5 min, incubated with 1ml MB#1 buffer (10 mM Tris-HCl, pH 7.5, 50 mM NaCl, 5 mM MgCl2, 1 mM CaCl2, 0.2% NP-40, 1x Roche cOmplete EDTA-free (Roche diagnostics, 04693132001)) and washed twice with MB#1 buffer. MNase concentration was predetermined using MNase titration experiments exploring 2.5-20U of MNase per million cells. We selected the 5U/million cells MNase concentration that gave us ∼90% mononucleosomes. Chromatin was digested with MNase for 10 min at 37 °C and digestion was stopped by adding 8 ul of 500 mM EGTA and incubating at 65 °C for 10 min.

Following dephosphorylation with rSAP (NEB #M0371) and end polishing using T4 PNK (NEB #M0201), DNA polymerase Klenow fragment (NEB #M0210) and biotinylated dATP and dCTP (Jena Bioscience #NU-835-BIO14-S and #NU-809-BIOX-S, respectively), ligation was performed in a final volume of 2.5 ml for 3h at room temperature using T4 DNA ligase (NEB #M0202). Dangling ends were removed by a 5 min incubation with Exonuclease III (NEB #0206) at 37 °C and biotin enrichment was done using 20 ul DynabeadsTM MyOneTM Streptavidin C1 beads (Invitrogen #65001). Libraries were prepared with the NEBNext Ultra II Library Preparation Kit (NEB #E7103). Samples were sequenced on a combination of Illumina’s NovaSeq 6000 and HiSeq 2500 at Novogene.

#### Contact enrichment analysis

Micro-C contacts that were mapped in one end to a 5kb window around the centers of dREG sites that overlapped with JIA risk associated SNPs and on the other end to a 5kb window around the transcription start site (TSS) of genes within 1Mbp from the center were captured. We used a LOWESS regression-based contact caller(14) to estimate enrichment of contacts relative to the expected given the contacts-by-distance decay in regions of up to 1Mbp from the center of the dREG peak – at the TSS orientation. We used Fisher’s exact p-values corrected by Benjamini-Hochberg false discovery rate (BH FDR) of 0.1 as a cutoff for a significant interaction between the JIA risk-associated SNP overlapping dREG site and a gene TSS.

## Results

### JIA SNPs within monocyte-macrophages dREG peaks

Genotyping data from children with JIA and healthy controls was analyzed to identify SNPs within candidate regulatory regions, identified by open chromatin and within H3K4me1/H3K27ac peaks, that were more common in children with JIA than the controls (p< 0.05) as described in the *Methods* section. Using bedtools intersect, we intersected the JIA SNPs (n=846) with dREG GRO-seq peaks in both monocytes and macrophages to identify candidate SNPs within regulatory regions. We identified 190 SNPs within dREG peaks in monocytes and 126 SNPs in macrophages. Of these, 101 SNPs were common between monocytes and macrophages (**Figure 1)**.

**Figure 1.**
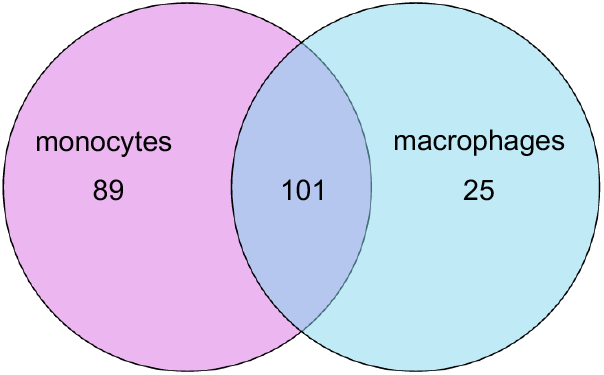
Venn diagram showing the numbers of candidate causal SNPs within dREG peaks identified in monocytes (n=89), macrophages (n=25) or common to both monocytes and macrophages (n=101).

The dREG peaks containing particular SNPs in monocytes and macrophages are shown in **Supplemental Tables 2** and **Supplemental Table 3**. Interestingly, the *C5orf56/IRF1* locus contained 27 and 20 SNPs in monocytes and macrophages respectively. Within the known PRM1-RM12 JIA risk haplotype region analyzed by Crinzi et al(8), 51 and 18 SNPs were located in this risk region, indicating an importance in these haplotypes regarding JIA genetic risk.

### Micro C data

We used MicroC data to identify gene promoters interacting with the regulatory regions harboring the candidate SNPs. MicroC data in monocytes identified multiple genes/transcripts whose promoters interact with the enhancers harboring the SNPs of interest, identifying likely target genes. The list of genes contacting dREG peaks is provided in **Table 2**, Micro-C data analysis of the *TYK2-ICAM1* region is shown in **Figure 2**.

**Table 2:** Genes/Transcripts Interacting With dREG-GROseq Peaks That Harbor Candidate JIA Causal Variants Identified by MicroC In Monocytes.

| Genes | ncRNAs | Transcripts-Unassigned Function |
| --- | --- | --- |
| CBLL1 | LINC01359 | AC002467.1 |
| ICAM1 | LINC02863 | AC009121.3 |
| ICAM5 |  | AC009121.4 |
| IRF1-AS1 |  | AC011511.4 |
| P4HA2 |  | AC116366.3 |
| RSBN1 |  | AL137856.1 |
| SLC22A4 |  | AL357078.2 |
| SOCS1 |  |  |
| TNP2 |  |  |
| TYK2 |  |  |
| ZGLP1 |  |  |

**Figure 2.**
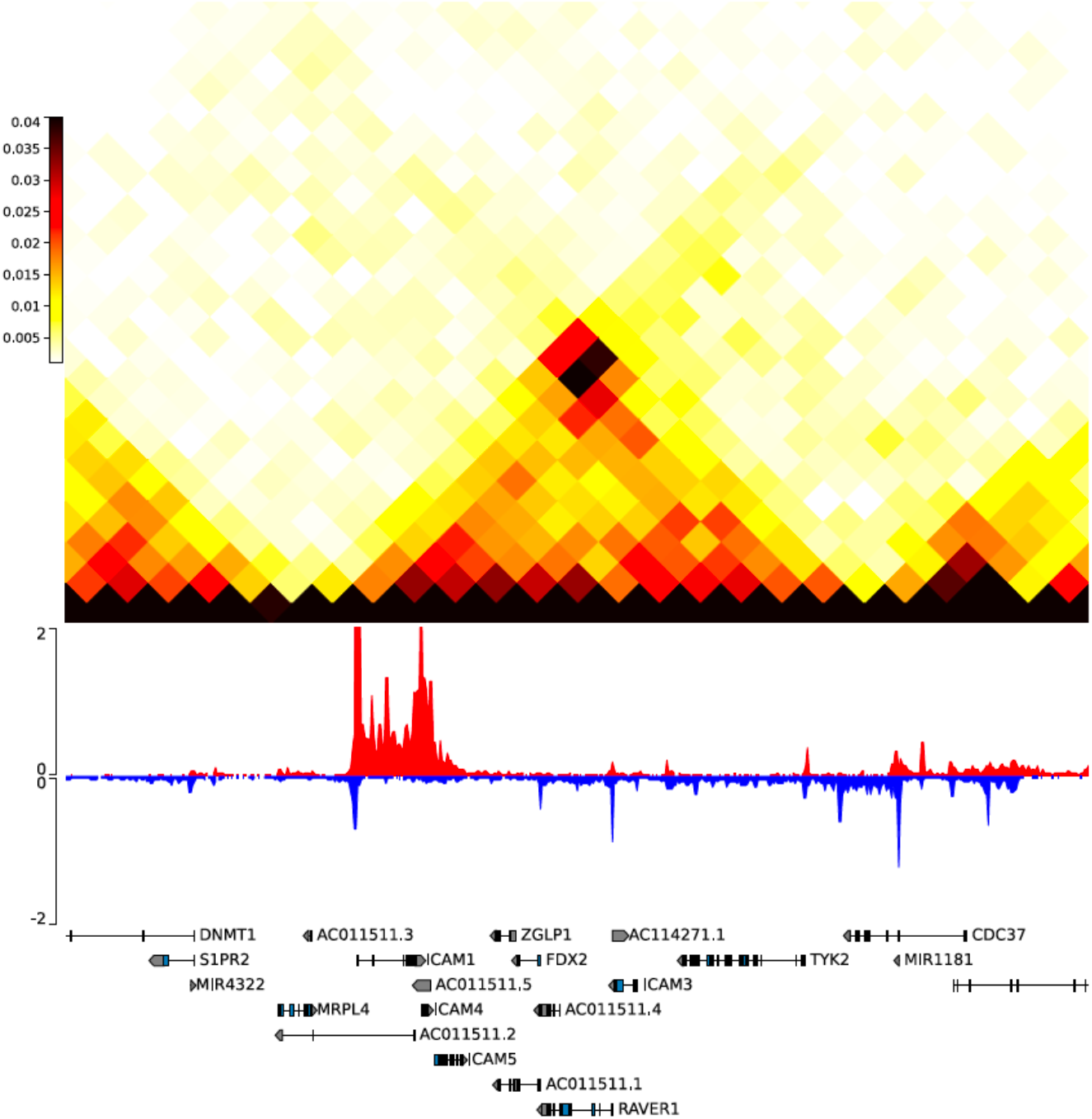
Heat map derived from MicroC analysis of resting human CD14+ monocytes. Shown are the regions encompassing ICAM1. RNA PolII positioning, as detected by GROseq, is shown on the sense (red) and antisense (blue) strands. Note that chromatin shows regions of contact forming micro-compartments (black and red triangles) that interact with the ICAM1 promoter. These findings are consistent with the presence of internal enhancers that regulate these genes.

Many of these genes are of direct interest to the pathobiology of JIA, which is characterized by exaggerated leukocyte responses to interferon gamma(15) and altered expression of multiple interferon-regulated genes(16). For example, SOCS1 is a known regulator of TLR-induced interferon pathways in monocytes(17), and TYK2 is required for normal interferon responses *in vivo*(18).

It is important to note that physical interaction, e.g, between a promoter and an enhancer, is not prima facie evidence that a regulatory relationship exists between those genomic elements(19).

Such relationships must be shown experimentally, e.g., with CRISPRi(20), where enhancer function can be attenuated and expression levels of candidate enhancer targets queried using quantitative rtPCR.

## Discussion

This study demonstrates how, using existing genotyping data and dREG, one can refine the functional regions likely to harbor SNPs that drive risk for JIA and other autoimmune diseases where risk is driven by the non-coding genome(9). Furthermore, using MicroC to visualize the 3D genome at high resolution, one can significantly narrow the search for target genes (i.e., the genes whose expression levels are influenced by the risk-driving variants).

In a previous study, we used Juicebox software analysis of HiC data to interrogate the JIA risk regions and identify the ”genomic space” in which JIA risk is likely to be exerted, and identified 237 genes that were within the same topologically-associated domains as putative enhancers (H3K27ac-ChIPseq peaks) within the JIA risk loci (10). Hi-C is a widely method for studying genome-wide chromatin interactions(21). Hi-C is sensitive for determining megabase-scale interactions and is particularly useful for determining large-scale chromatin structure(22). However, since Hi-C uses restriction enzymes, the coverage of Hi-C is not comprehensive and instead is fragmented. Furthermore, with Hi-C, the average chromatin fragment size is 4 kb, which is larger than most regulatory elements. Compared to Hi-C, Micro-C identifies interactions at a resolution of 200 bp, including interactions around structural variants(23). Furthermore, Micro-C is capable of capturing long-range chromatin interactions while maintaining its high resolution, identifying physical interactions between distal regulatory elements and promoters(10, 24). The precision and sensitivity of MicroC analyses narrows the list of candidate target genes and transcripts (in monocytes, at least) to 20.

A curious finding in this study is the Micro-C identified interactions between PROseq-dREG identified peaks and non-coding transcripts, which we have not observed in other immune cells (unpublished observations). Long non-coding RNAs (lncRNAs) are known to recruit regulatory complexes and modulate the expression of nearby genes, adding additional fine-tuning of gene expression at the local level(25). Our findings, however, must be interpreted cautiously. Some of these transcripts may fall in the same regulatory interaction domain as a nearby protein coding gene and the actual target is the protein coding gene.

We have previously postulated that genetic variants of JIA may confer disease risk by modifying both the innate and adaptive arms of the immune system. The involvement of the adaptive immune system is supported by the fact that the Class II HLA locus confers the strongest risk for the disease(5). At the cellular level, there is a demonstrated imbalance between Tregs and autoreactive Th1/Th17 cells in patients with oligoarticular and polyarticular JIA(26). However, neither the presence of autoantibodies (which are relatively scarce in JIA and do not distinguish children with the disease from healthy children(27)) nor of autoreactive lymphocytes fully account for the pathogenesis of many autoimmune diseases without the participation of innate immunity(28, 29).

Monocytes and macrophages are key participants in the innate immune system and are critical in coordinating the initiation, recruitment, and resolution stages of many autoimmune diseases(8). In addition, they display many immunoregulatory effects that impact inflammation that regulate tissue repair and regeneration. Schmidt et al. identified a distinct activation pattern in synovial fluid monocytes and functional alterations with inflammatory and regulatory features, such as an increased ability to induce T-cell activation and resistance to cytokine production, indicating an important role 5for the innate immune system activation in oligoarticular JIA(13, 30). Nonclassical CD14+CD16++ macrophages and intermediate CD14++CD16+ macrophages account for a smaller proportion of circulating macrophages, comparatively to classical macrophages, and can expand substantially in inflammatory environments(12). While no significant increase in classical macrophages has been observed in patients with polyarticular JIA, the frequency of intermediate macrophages is expanded(31). Raggi et al. determined that synovial macrophages in the hypoxic inflamed joints of children with oligoarticular JIA overexpressed triggering receptors expressed on myeloid cells (TREM)-1 and released VEGF and osteopontin(32). TREM-1 reverses the immunoregulatory effect of hypoxia and drives proinflammatory reprogramming whereas VEGF and osteopontin drive neoformation of blood vessels by enhancing epithelial cell survival/proliferation, as well as monocyte cell recruitment/activation. However, as with most studies of JIA synovial fluid cells, it is impossible to determine whether these findings represent immune perturbations characteristic of the disease state or are the predictable consequences of chronic tissue inflammation.

In conclusion, we have shown that the use of genotyping data, measures to identify active transcriptional regulatory elements, and high-dimensional 3D chromatin mapping can considerably reduce the 3 dimensional chromatin space to be explored to define genetic mechanisms that contribute to risk in JIA. While further experimental evidence is required to finally determine that a candidate SNP is truly risk-driving, and that a candidate target gene is affected by the SNP(s) of interest, the approach we describe here provides an effective method for focusing that additional experimental work.

## Author Contributions

EH – Assisted in designing the study, performed BedTools analyses of GROseq data, and assisted in writing the manuscript.

GB – Performed MicroC and GROseq analyses and prepared the figures for the manuscript. AH – Assisted in analysis of MicroC data

ER – Performed GROseq experiments

EC – Provided assistance with BedTools analysis of GROseq data KJ – Assisted in data interpretation and preparation of the manuscript

CDG – Supervised PROseq and MicroC procedures and analyses, assisted in data interpretation and preparation of the manuscript.

JNJ - Designed the study, supervised its implementation, and assisted in data interpretation and preparation of the manuscript.

## Acknowledgments

The authors wish to thank Drs. Susan Thompson and Marc Sudman at the Cincinnati Children’s Medical Center (CCHMC) for access to CCHMC genotyping data.

## Data Availability Statement

The datasets for this study can be found in the Gene Expression Omnibus (GEO), accession numbers GSM1183908 and GSM1183912. MicroC and raw GRO-seq data are been posted on dbGaP under accession number phs002146.v1.p1.

## References

1. Gortmaker SL, Sappenfield W. Chronic childhood disorders: prevalence and impact. Pediatr Clin North Am. 1984;31(1):3–18.

2. Singsen BH. Rheumatic diseases of childhood. Rheum Dis Clin North Am. 1990;16(3):581–99.

3. Prahalad S, Zeft AS, Pimentel R, Clifford B, McNally B, Mineau GP, et al. Quantification of the familial contribution to juvenile idiopathic arthritis. Arthritis Rheum. 2010;62(8):2525–9.

4. McIntosh LA, Marion MC, Sudman M, Comeau ME, Becker ML, Bohnsack JF, et al. Genome-Wide Association Meta-Analysis Reveals Novel Juvenile Idiopathic Arthritis Susceptibility Loci. Arthritis Rheumatol. 2017;69(11):2222–32.

5. Hinks A, Cobb J, Marion MC, Prahalad S, Sudman M, Bowes J, et al. Dense genotyping of immune-related disease regions identifies 14 new susceptibility loci for juvenile idiopathic arthritis. Nat Genet. 2013;45(6):664–9.

6. Jiang K, Zhu L, Buck MJ, Chen Y, Carrier B, Liu T, et al. Disease-Associated Single-Nucleotide Polymorphisms From Noncoding Regions in Juvenile Idiopathic Arthritis Are Located Within or Adjacent to Functional Genomic Elements of Human Neutrophils and CD4+ T Cells. Arthritis Rheumatol. 2015;67(7):1966–77.

7. Zhu L, Jiang K, Webber K, Wong L, Liu T, Chen Y, et al. Chromatin landscapes and genetic risk for juvenile idiopathic arthritis. Arthritis Res Ther. 2017;19(1):57.

8. Crinzi EA, Haley EK, Poppenberg KE, Jiang K, Tutino VM, Jarvis JN. Analysis of chromatin data supports a role for CD14+ monocytes/macrophages in mediating genetic risk for juvenile idiopathic arthritis. Frontiers in Immunology. 2022;13.

9. Farh KK, Marson A, Zhu J, Kleinewietfeld M, Housley WJ, Beik S, et al. Genetic and epigenetic fine mapping of causal autoimmune disease variants. Nature. 2015;518(7539):337–43.

10. Jiang K, Kessler H, Park Y, Sudman M, Thompson SD, Jarvis JN. Broadening our understanding of the genetics of Juvenile Idiopathic Arthritis (JIA): Interrogation of three dimensional chromatin structures and genetic regulatory elements within JIA-associated risk loci. PLoS One. 2020;15(7):e0235857.

11. Kim-Hellmuth S, Aguet F, Oliva M, Munoz-Aguirre M, Kasela S, Wucher V, et al. Cell typespecific genetic regulation of gene expression across human tissues. Science. 2020;369(6509).

12. Wu C-Y, Yang H-Y, Huang J-L, Lai J-H. Signals and Mechanisms Regulating Monocyte and Macrophage Activation in the Pathogenesis of Juvenile Idiopathic Arthritis. International Journal of Molecular Sciences. 2021;22(15):7960.

13. Schmidt T, Dahlberg A, Berthold E, Król P, Arve-Butler S, Rydén E, et al. Synovial monocytes contribute to chronic inflammation in childhood-onset arthritis via IL-6/STAT signalling and cell-cell interactions. Frontiers in Immunology. 2023;14.

14. Chu T, Rice EJ, Booth GT, Salamanca HH, Wang Z, Core LJ, et al. Chromatin run-on and sequencing maps the transcriptional regulatory landscape of glioblastoma multiforme. Nat Genet. 2018;50(11):1553–64.

15. Throm AA, Moncrieffe H, Orandi AB, Pingel JT, Geurs TL, Miller HL, et al. Identification of enhanced IFN-γ signaling in polyarticular juvenile idiopathic arthritis with mass cytometry. JCI Insight. 2018;3(15).

16. Knowlton N, Jiang K, Frank MB, Aggarwal A, Wallace C, McKee R, et al. The meaning of clinical remission in polyarticular juvenile idiopathic arthritis: gene expression profiling in peripheral blood mononuclear cells identifies distinct disease states. Arthritis Rheum. 2009;60(3):892–900.

17. Prele CM, Woodward EA, Bisley J, Keith-Magee A, Nicholson SE, Hart PH. SOCS1 regulates the IFN but not NFkappaB pathway in TLR-stimulated human monocytes and macrophages. J Immunol. 2008;181(11):8018–26.

18. Prchal-Murphy M, Semper C, Lassnig C, Wallner B, Gausterer C, Teppner-Klymiuk I, et al. TYK2 kinase activity is required for functional type I interferon responses in vivo. PLoS One. 2012;7(6):e39141.

19. Tarbell E, Jiang K, Hennon TR, Holmes L, Williams S, Fu Y, et al. CD4+ T cells from children with active juvenile idiopathic arthritis show altered chromatin features associated with transcriptional abnormalities. Sci Rep. 2021;11(1):4011.

20. Jiang K, Liu T, Kales S, Tewhey R, Kim D, Park Y, et al. A systematic strategy for identifying causal single nucleotide polymorphisms and their target genes on Juvenile arthritis risk haplotypes. BMC Med Genomics. 2024;17(1):185.

21. Lee BH, Wu Z, Rhie SK. Characterizing chromatin interactions of regulatory elements and nucleosome positions, using Hi-C, Micro-C, and promoter capture Micro-C. Epigenetics Chromatin. 2022;15(1):41.

22. Davies JOJ, Oudelaar AM, Higgs DR, Hughes JR. How best to identify chromosomal interactions: a comparison of approaches. Nature Methods. 2017;14(2):125–34.

23. Hansen AS, Hsieh TS, Cattoglio C, Pustova I, Saldaña-Meyer R, Reinberg D, et al. Distinct Classes of Chromatin Loops Revealed by Deletion of an RNA-Binding Region in CTCF. Mol Cell. 2019;76(3):395-411.e13.

24. Hsieh T-HS, Cattoglio C, Slobodyanyuk E, Hansen AS, Rando OJ, Tjian R, et al. Resolving the 3D Landscape of Transcription-Linked Mammalian Chromatin Folding. Molecular Cell. 2020;78(3):539-53.e8.

25. Engreitz JM, Haines JE, Perez EM, Munson G, Chen J, Kane M, et al. Local regulation of gene expression by lncRNA promoters, transcription and splicing. Nature. 2016;539(7629):452–5.

26. Lin YT, Wang CT, Gershwin ME, Chiang BL. The pathogenesis of oligoarticular/polyarticular vs systemic juvenile idiopathic arthritis. Autoimmun Rev. 2011;10(8):482–9.

27. McGhee JL, Kickingbird LM, Jarvis JN. Clinical utility of antinuclear antibody tests in children. BMC Pediatr. 2004;4:13.

28. Ma WT, Gao F, Gu K, Chen DK. The Role of Monocytes and Macrophages in Autoimmune Diseases: A Comprehensive Review. Front Immunol. 2019;10:1140.

29. Laria A, Lurati A, Marrazza M, Mazzocchi D, Re KA, Scarpellini M. The macrophages in rheumatic diseases. J Inflamm Res. 2016;9:1–11.

30. Schmidt T, Berthold E, Arve-Butler S, Gullstrand B, Mossberg A, Kahn F, et al. Children with oligoarticular juvenile idiopathic arthritis have skewed synovial monocyte polarization pattern with functional impairment-a distinct inflammatory pattern for oligoarticular juvenile arthritis. Arthritis Res Ther. 2020;22(1):186.

31. Gaur P, Myles A, Misra R, Aggarwal A. Intermediate monocytes are increased in enthesitis-related arthritis, a category of juvenile idiopathic arthritis. Clin Exp Immunol. 2017;187(2):234–41.

32. Raggi F, Pelassa S, Pierobon D, Penco F, Gattorno M, Novelli F, et al. Regulation of Human Macrophage M1-M2 Polarization Balance by Hypoxia and the Triggering Receptor Expressed on Myeloid Cells-1. Front Immunol. 2017;8:1097.

